# Human postmortem studies reveal tissue-specific differences amongst TB-patient groups

**DOI:** 10.1101/2023.03.14.532701

**Authors:** Gift Ahimbisibwe, Marjorie Nakibuule, Marvin Martin Ssejoba, David Oyamo, Rose Mulwana, Josephine Nabulime, Febronius Babirye, Abdusalaamu Kizito, Herve Lekuya, Akello Suzan Adakun, Robert Lukande, Andrew Kyazze, Irene Andia Biraro, Stephen Cose

## Abstract

If we are to break new grounds in TB research, we need to have a complete understanding of what is occurring at the site of infection in humans. Postmortem studies give us an opportunity to compare TB-involved and -uninvolved tissues, in both diseased and non-diseased individuals. We examined the feasibility of carrying out a postmortem study in Mulago and Kiruddu National Referral Hospitals in Uganda, to determine whether we could use immune cells collected postmortem for immunological studies. We report that we can consent the Next-of-Kin, perform postmortem procedures and process tissues within 8 hours of death, and that immune cells remain viable and functional up to 14 hours after death. We found subtle differences in T cell subsets within TB groups. We found a depletion of the CD4 CD69^+^CD103^+^ T cell subset in the lungs and BAL, which was associated with HIV, and that the CD8 CD69^+^CD103^-^ T cell subset was depleted in BAL only, and was associated with TB. Our data show overall changes Tissue Resident Memory T cells within, and between, TB-infected and TB-uninfected human lungs.

**Summary:**

1. Coroner led postmortem studies are possible in Uganda, samples processed within 8 hours from death
2. Cells from samples collected postmortem are viable and functional
3. HIV associated depletion of CD4 CD69^+^/CD103^+^ T cell subset in lungs and BAL
4. CD8 CD69^+^/CD103^-^ depletion in BAL associated with TB

## Introduction

There is no question that the SARS-CoV-2 (COVID-19) pandemic has reversed many of the gains made in fighting the effort to control the spread of tuberculosis (TB), and has set back the fight against TB by years (Jeremiah et al., 2022). Nevertheless, we cannot blame our lack of success in controlling TB on COVID-19 alone. We have lived with TB long enough not to have it as the leading cause of death from a single infectious disease prior to SARS-CoV-2, and by all counts set to regain its number one spot again (WHO, 2022). The major challenges hindering us from hitting the global 2030 End TB reduction targets range from a lack of efficient TB vaccines, diagnostic challenges, drug resistance and inaccessibility to effective chemotherapy (Dye et al., 2013; WHO, 2022).

The best intervention against TB, or in fact any infectious disease, is one that breaks the transmission cycle. Lessons from other infectious diseases that have now been eradicated, such as smallpox and rinderpest, have shown that vaccination is the best way to break transmission; polio is almost eradicated as well due to the WHO’s determined vaccination strategy (WHO, 2021). TB vaccination has been extensively taken up by most TB endemic countries. However, it is well-known that BCG, currently the only licensed TB vaccine, isn’t effective against the most transmissible form of the disease in endemic countries: adult pulmonary TB (Martinez et al., 2022). We need to develop more effective TB vaccines and develop shorter drug regimens, amongst a plethora of other interventions (WHO, 2022). The last few years have witnessed a reawakening of novel TB vaccine approaches. There is hope in some current trial vaccines (M72, BCG REVAC, VPM1002, H56/IC31) but tertiary trial results will not be out for several years (Atulomah et al., 2022; Frick, 2021). In the meantime, more needs to be done. We need to come up with vaccine strategies based on deeper insights into the immunity underlying TB, and not just in blood, but most importantly at the site of infection in humans – otherwise we are simply designing our next vaccine empirically.

TB disease typically affects the lungs in humans (Pulmonary TB), with an ability to disseminate and affect other tissue sites. It is now widely known that tissue based immune cells are present within the lungs to fight off lung specific infections such as TB and influenza (Beura et al., 2019; Mueller et al., 2013). The human lung is a highly vascularised tissue and cells within the lung are located either within the parenchyma itself, or embedded within the vasculature (Anderson et al., 2014). The cells located in the lung vasculature are believed to be in circulation and capable of moving back to the general body circulation, whereas those within the parenchyma are believed to be a sessile, Tissue Resident Memory (TRM) population (Mueller et al., 2013; Schenkel and Masopust, 2014; Snyder et al., 2019).

Lung TRMs express distinct tissue retention and survival markers including CD69, CD103, CD101, CD49a, CD49d and CXCR3, said to be a result of tissue specific cues and antigens within the lung microenvironment (Thome and Farber, 2015). The positioning of these cells within the tissues is strategic and enables a rapid and immediate response upon subsequent antigen encounter. More importantly, these cells have been shown to provide more intensive antigen clearance compared to their non TRM counterparts, which is attributed to higher rates of cytokine production and proliferation in response to the presence of antigen (Gray et al., 2018). Indeed, adoptive transfer studies, in both murine and NHP models, show that CXCR3 and CD69/CD103 positive populations were found to be better at *M.tb* clearance compared to their negative counterparts (Corleis et al., 2019; Perdomo et al., 2016). On the other hand, cells within the lung vasculature respond to antigen by crossing to the parenchyma. Cells that express the lung homing receptor CXCR3 were found to provide superior protection against TB because of their ability to get to the site of infection (Sakai et al., 2014). It is therefore important to phenotype and understand cell composition and roles within the lungs in TB disease and health if we are to exploit the cellular immune system in developing novel vaccine strategies.

Overall, animal models have been used extensively to study and characterize TRMs (Fonseca et al., 2017). However, the inability to mimic the complex natural history of TB disease progression, and perhaps more importantly, reactivation tuberculosis commonly seen in adults, means that the relevant immune response may be overlooked in animal models (Bucsan et al., 2019; Zhan et al., 2017). This highlights the importance of studying TRMs in the context of TB disease, and specifically within human tissues. The study of TB in human tissues was last extensively done before the advent of effective chemotherapy (listed, 1854; Rozenblat, 1949; Saye, 1954). Unfortunately, now with major technological advances, it is increasingly difficult to get human tissues. As a result, human TB immunology research is extensively focused on peripheral blood, whereas tissue immunology studies rely heavily on animal models which do not develop the disease as it exists in humans. There have been efforts to obtain and study human tissues by lung resection, but the issue with these studies is that the tissues obtained are from critically ill TB patients undergoing surgical resection due to severe lung complications (Ardain et al., 2019; Karimi et al., 2014; Ogongo et al., 2020). Under such circumstances, it is difficult to get “healthy” tissue for comparison, either from the same individual or another healthy individual. This has hampered the field of human lung immunology for many years.

One approach to help advance the field is to undertake postmortem studies to better understand the disease at the site of infection. Postmortem studies were extensively performed right up into the 1950’s but have subsequently fallen out of favour (Hunter et al., 2009; Hunter, 2016). There are a number of reasons for this, ranging from difficulty in obtaining consent (Loughrey et al., 2000), concern over misdiagnosis at death (Royal College of Pathologists of Australasia Autopsy Working, 2004), physician reluctance (Sinard, 2001; Turnbull et al., 2015), age of patient at death (Hoyert, 2011; Sinard, 2001), cost (Britton, 1993), and scandal (Burton and Underwood, 2003). These issues may also be relevant in our setting in Uganda, although few studies have assessed such obstacles in any systematic way. This notwithstanding, in this study we show that we can undertake successful postmortems for medical research, and that human postmortem procedures remain a valuable and essential tool to both establish cause of death, and to advance our medical and scientific understanding (Loughrey et al., 2000).

We conducted a postmortem study to evaluate the feasibility of undertaking medical research in Mulago and Kiruddu National Referral Hospitals in Kampala, Uganda, to determine whether we could use immune cells collected postmortem for immunological studies. Postmortem studies can inform us, on a much greater scale than previously understood, and with new techniques now available, exactly how the immune system responds at the site of TB infection, in both health and disease. In this study we show that we can consent the next of kin(s) (NOKs), perform a full postmortem, collect and process tissues within 14 hours from death, and with no loss in cell viability or function. In studies such as this, we can directly compare involved and uninvolved tissues, and diseased and non-diseased tissues within the same individual. We show that we can get tissues from a variety of TB infectious and treatment states, as well as uninfected cases.

## Materials and Methods

### Ethics

This study obtained ethical approval from five ethics bodies: the Makerere University School of Biomedical Sciences Research & Ethics Committee, the Mulago National Referral Hospital Ethics Committee, the Kiruddu National Referral Hospital Ethics Committee, the Uganda National Council of Science and Technology Ethics Committee and the London School of Hygiene and Tropical Medicine Ethics Committee.

### Consent

Trained grief councillors were employed to sensitively obtain informed consent from the NOK as described by Uganda Law. The NOK gave consent for a full postmortem, for donation of tissues for medical research and for access to previous medical records. A case record form was used by the councillors to collect reasons for consent or decline from the NOK.

### Participants

We recruited deceased subjects from Mulago and Kiruddu National Referral Hospitals in Kampala, Uganda, between January 2021 and June 2022. Active TB subjects were recruited from Wards 4, 5 and 6 of Mulago National Referral Hospital (MNRH), plus the Infectious Diseases Ward at Kiruddu National Referral Hospital (KNRH). We used road traffic accident victims as our non-TB subjects, which were recruited from the Accident and Emergency Unit of MNRH (referred to as the Surgical Emergency Unit). Recruited subjects from this Unit were otherwise healthy at the time of death, with no obvious underlying morbidity or chest involvement at time of death. Patient demographics are described in Table 1.

**Table 1.** Patient demographics. Data shows collated patient demographics across the whole study.

|  | **S&E<sup>a</sup>** | **TB<sup>b</sup>** |
| --- | --- | --- |
| **Gender** |  |  |
| Male | 37 | 20 |
| Female | 10 | 13 |
| **Age Range** |  |  |
| 18-35 | 26 (26.3)<sup>c</sup> | 13 (28.6) |
| 36-55 | 12 (41.7) | 15 (43.8) |
| ≥56 | 9 (69.4) | 5 (67) |
| **HIV status** |  |  |
| Positive | 6 | 23 |
| Negative | 41 | 10 |
| **Cause of death** |  |  |
| *Trauma* |  |  |
| Head trauma | 35 |  |
| Abdominal | 8 |  |
| Gunshot | 1 |  |
| *Non trauma* |  |  |
| Intestinal obstruction | 2 |  |
| Duodenal ulcers | 1 |  |
| *TB Cases* |  |  |
| Pulmonary TB |  | 12 |
| Extrapulmonary TB |  | 13 |
| HIV associated |  | 8 |
| **Total** | 47<br>per group | 33 |

<sup>a</sup>Surgical and Emergency ward, <sup>b</sup>Tuberculosis Wards, <sup>c</sup>Parentheses represent average age

### Tissues and sample collection

A full postmortem was performed by the study pathologist on all study subjects to establish the cause of death and identify any underlying conditions that may have been missed during routine clinical and laboratory examination. Tissue samples, including lung, lung draining lymph nodes (Hilar Lymph Nodes - HLNs), spleen, distal lymph nodes (Iliac and inguinal lymph nodes – DLNs) and blood were collected. The solid tissues were placed in 50ml tubes with 20% FBS in RPMI medium. Bronchoalveolar lavage (BAL) washing of both the left and right lungs were performed with PBS to obtain BAL samples. Arterial blood was collected from the carotid artery into heparin tubes; arteries were considered instead of veins due to regular venous collapse after death. Sample tubes were tightly capped, placed in a rack in a sealable cool box and transported to the BSL3 laboratory at the MRC/UVRI and LSHTM Uganda Research Unit at room temperature. All sample processing was performed under BSL3 conditions.

### PBMC isolation

Heparinised blood was diluted with RPMI media and layered manually on FicollPaque PLUS media. PBMCs were then separated from the diluted heparinised by density centrifugation. The buffy coat was harvested and centrifuged to obtain a cell pellet. Red blood cells in the pellet were then lysed, the cells washed and then counted on an automated cell counter (TC20; Biorad).

### Cell isolation from tissues

#### 1. Lung

Cells were obtained from lung tissue by enzymatic digestion using collagenase D (1mg/ml) and DNase I (1g/ml) (hereafter referred to as “the enzyme mixture”) and physical disintegration using the gentleMACS Octo Dissociator (Milteyni Biotech) as described previously (Sathaliyawala et al., 2013). Briefly, the lungs were chopped into small pieces using fine scissors and forceps in a sterile petri dish, placed in the gentleMACS purple C tubes with the enzyme mixture, loaded on the gentleMACS Octo Dissociator and run-on Lung Program 1. The tubes were then incubated in a CO_2_ incubator for 25 minutes followed by being run on Lung Program 2 on the gentleMACS. The sample was then filtered through a 70 μm filter followed by a 40 μm one, centrifuged at 600 rcf for 5 minutes to obtain a cell pellet. Residual red blood cells were lysed using ACK lysis buffer, leaving behind a pure black cell pellet. We assume the cell pellet from the draining lungs was black due to a lifetime exposure to carbon residues as a result of inhalation of carbon fumes from vehicles, firewood, and charcoal smoke. The cells were washed in RPMI and counted using an automated TC20 cell counter (Biorad)

#### 2. Lymph nodes

Lymph nodes were first cleaned by teasing the lymph node tissue from surrounding fat using forceps and scissors, chopped into small pieces using fine scissors and forceps in a sterile petri dish and mixed with RPMI media. The resultant mixture was then filtered through a 70 μm filter followed by a 40 μm filter, then centrifuged at 600 rcf for 5 minutes to obtain a cell pellet. Red blood cells were lysed using ACK lysis buffer. The resultant pellet was washed with RPMI and counted using the automated TC20 cell counter.

#### 3. Spleen

Spleen sections were chopped into small pieces using fine scissors and forceps in a sterile petri dish and mixed with RPMI media. The resultant mixture was then filtered through a 70 μm filter and centrifuged at 600 rcf for 5 minutes to obtain a cell pellet. The pellet was then reconstituted with RPMI and layered manually onto FicollPaque PLUS media. Spleen mononuclear cells were then obtained by density centrifugation. The buffy coat was harvested and centrifuged to obtain a cell pellet. Residual red blood cells in the pellet were lysed using ACK lysis buffer, and the cells then washed and counted using an automated TC20 cell counter.

#### 4. Bronchoalveolar lavage fluid

BAL fluid was filtered through a 70 μm filter followed by a 40 μm filter, then centrifuged at 600 rcf for 5 minutes to obtain a cell pellet. Red blood cells were lysed using ACK lysis buffer. The resultant pellet was washed with RPMI and counted using an automated TC20 cell counter.

### T-SPOT®.TB assay

This assay was performed using the T-SPOT.TB kit (TB.300, Oxford Immunotec) to enumerate individual TB-specific activated effector T cells producing IFN-γ. Samples were run in duplicate. The procedure was performed as per insert, except for incubation time (36 hrs) and media (RPMI plus 10% FBS). Extensive testing revealed that this time and media yielded best results from our deceased subjects, above that of the recommended protocol for live venous blood. Resultant spots were read using an ELISPOT reader (AID *i*Spot ELR08IFL). Results were interpreted as per insert.

### Cell phenotypic analysis by flow cytometry

A 29-colour antibody T and B cell panel was used to phenotype the cells isolated from each of the tissues. The gating strategy for PBMCs and tissues are shown in Figures 1 and 2, respectively. Antibodies used were CD3 (PACIFIC BLUE HIT3a BioLegend), CD38 (BV510 HIT2 Biolegend), CCR7 (BV605 G043H7 Biolegend), PD-1 (BV650 NAT105 Biolegend), CD69 (BV711 FN50 Biolegend), CD28 (BV750 CD28.2 Biolegend), CD27 (BV785 O323 Biolegend) CD8 (SPARK BLUE 550SK1 Biolegend) FCLR4 (PE413D12 Biolegend), CD127 (SPARK YG581 A019B5 Biolegend) CD4 (PECY5 RPA-T4 Biolegend), CD57 (PECY7 HNK-1 Biolegend), IgG (APC M1310G05 Biolegend), KLRG1 (ALEXAFLUOR 647 SA231A2 Biolegend), CD21 (ALEXAFLUOR 700 Bu32 Biolegend) HLADR (APC-FIRE 810 I243 Biolegend), LIVE/DEAD (ZOMBIE UV Biolegend) CD103 (BUV395 Ber ACT8 BD), CD25 (BUV496 M-A251 BD), CD196 (BUV563 11A9 BD), CD278 (BUV661 DX29 BD), IgM (BUV737 UCH-B1 BD), CD45RA (BUV805 HI100 BD), CD5 (BV421 UCHT2 BioLegend), CD24 (FITC eBioSN3 eBioscience), CD183 (BB700 1C6/CXCR3 BD), CD10 (PerCP efluor710 SN5c Life technologies), IgD (PE-Dazzle594 IA6-2 BD), CD19 (APC-H7 SJ25C1 BD),

**Figure 1.**
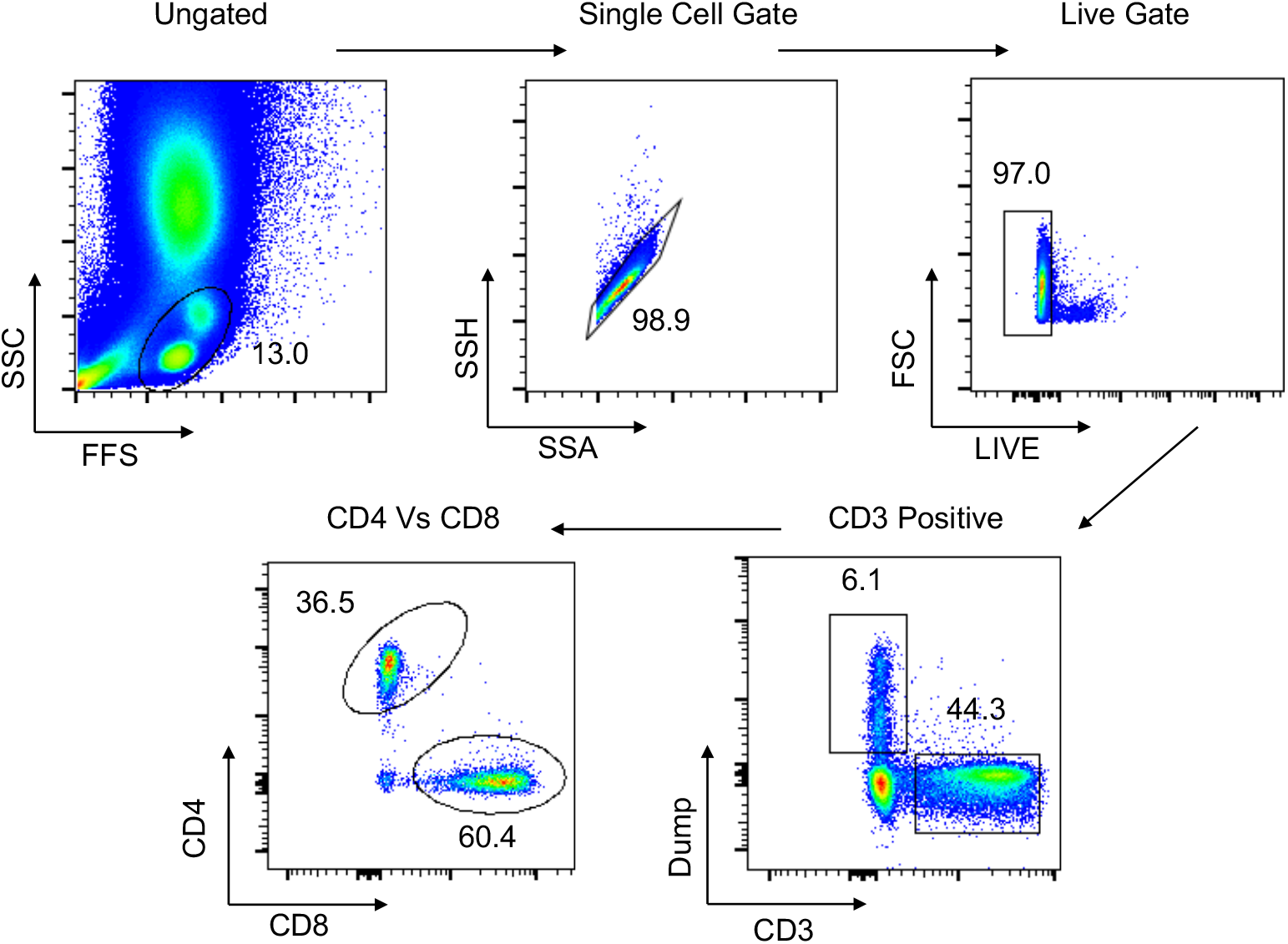
Gating strategy for flow cytometry in PBMCs. Representative plot of total cells from arterial PBMCs taken postmortem. Total cells are first gated by FSC/SSC, then singlets isolated, followed by selection of live cells and then CD3 positive cells. The dump gate, used to make the CD3 gate cleaner, consists of CD19/CD56/CD14. Finally, cells are separated based on CD4 and CD8, followed by specific staining as represented in the Results. Data comes from a 65-year-old male HIV-/TB-subject recruited from the SEU.

**Figure 2.**
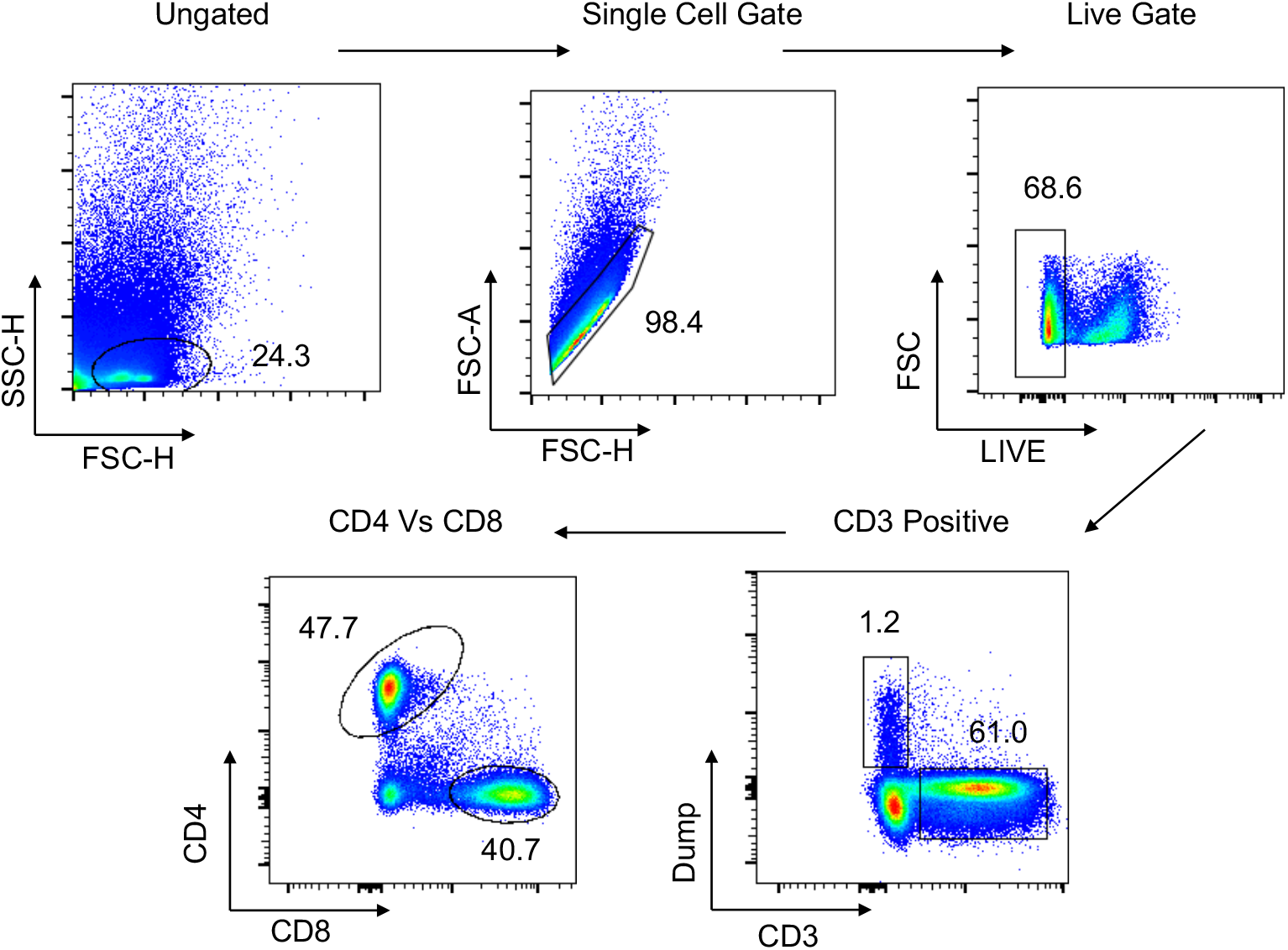
Gating strategy for flow cytometry in tissues. Representative plot of total cells from lung tissue taken postmortem. Total cells are first gated by FSC/SSC, then singlets isolated, followed by selection of live cells and then CD3 positive cells. The dump gate, used to make the CD3 gate cleaner, consists of CD19/CD56/CD14. Finally, cells are separated based on CD4 and CD8, followed by specific staining as represented in the Results. A similar gating strategy was employed for all other tissues. Data comes from a 65-year-old male HIV-/TB-subject recruited from the SEU.

### Phenotypic staining

Viability staining was performed first with the fixable viability dye, zombie UV, for 20 minutes in the dark at room temperature. Cells were washed and resuspended in 100 μl of surface antibody cocktail for 20 minutes in the dark at 4°C, after which they were washed to remove excess antibody. Cells were acquired using a Cytek Aurora Spectral Analyser. All flow cytometry data were analysed using FlowJo version 10.8.1. Compensation controls to remove spectral overlap and Fluorescence Minus One (FMO) were used to establish gates.

### Statistical Analysis

FlowJo and GraphPad prism version 9 software was used for graphical Representation and statistical analysis.

## Results

To determine whether tissue-specific immunology studies could be performed on tissues donated for medical research following death, we assessed two essential elements: 1) could we get consent from the next-of-kin (NoK), and if so, 2) how viable would cells be following death and the postmortem process to collect tissues for medical research.

We found that we could consent the NoK with very high consent rates on the Surgical and Emergency (S&E) wards (94% of those approached, Table 2), and more modest rates of consent on the TB Wards (35% of those approached, Table 2). The latter consent rate was in line with a previous study assessing consent across all Wards at Mulago National Referral Hospital (MNRH) (Cox et al., 2011). The high rate of consent on the S&E Ward was a surprise, as we had assumed a sudden, unexpected death might be more traumatic for the NOK, whereas the consent rate on the TB wards might be higher because the patient and family have usually built up a rapport with hospital staff and may therefore be more willing to consent into a postmortem study for their deceased relative. However, most NOKs believed that understanding the reason for a loved one’s sudden death helps them come to terms with their loss. The low consent rate was mainly attributed to religious norms that don’t agree with postmortems.

**Table 2.** Reasons for consent and decline. Whilst approaching families for consent into the study, we asked a short questionnaire on why they did or didn’t accept for their deceased family member to be a part of the study, and for the use of their tissues for medical research. Data shows the responses given in the families that agreed to answer the questionnaire.

| **Numbers** |  | **<sup>a</sup>S&E** | **TB Wards** |
| --- | --- | --- | --- |
| Met study criteria |  | 50 | 94 |
| Consented |  | 47 | 33 |
| Declined |  | 3 | 61 |
| Consent rate |  | 94% | 35% |
| **Reasons for Consent** |  | **S&E (47)** | **TB Wards (33)** |
| **Gender** | Male | 13 | 10 |
|  | Female | 34 | 23 |
| Knowing the cause of death will help prevent more deaths |  | 5 | 10 |
| It is important in settling NSSF, deceased's bank payments |  | 8 | 8 |
| Understanding the reason for a loved one's death helps them come to terms with their loss |  | 17 | 9 |
| It sounds like the right thing to do |  | 9 | 1 |
| It is a way of saving future lives |  | 4 | 5 |
| **Reasons for Decline** |  | **S&E (3)** | **TB Wards (61)** |
| **Gender** | Male | 1 | 27 |
|  | Female | 2 | 34 |
| Religious norms |  | 1 | 16 |
| Cultural norms |  | 1 | 12 |
| Fear of body disorientation |  | 1 | 8 |
| Process will delay funeral arrangements |  |  | 10 |
| PM not necessary |  |  | 2 |
| Lack of authority |  |  | 3 |
| Not sure deceased would approve |  |  | 4 |
| No reason |  |  | 6 |

<sup>a</sup> Surgical and Emergency Unit

Having gained consent, the body was taken for a full postmortem and tissues taken to assess the viability of cells over time, from the time of death to the beginning of sample processing in the laboratory (Figure 3). We found that cells were viable up to 14 hours post death, but that there was considerable variability in viability, depending on the tissue sample. Arterial blood maintained a high level of viability out to 14 hours post death, but other tissues examined were considerably lower; around the 40% area of viability. The only tissue to show a significant difference in viability and time from death was the distal lymph node (DLN; iliac and inguinal lymph nodes).

**Figure 3.**
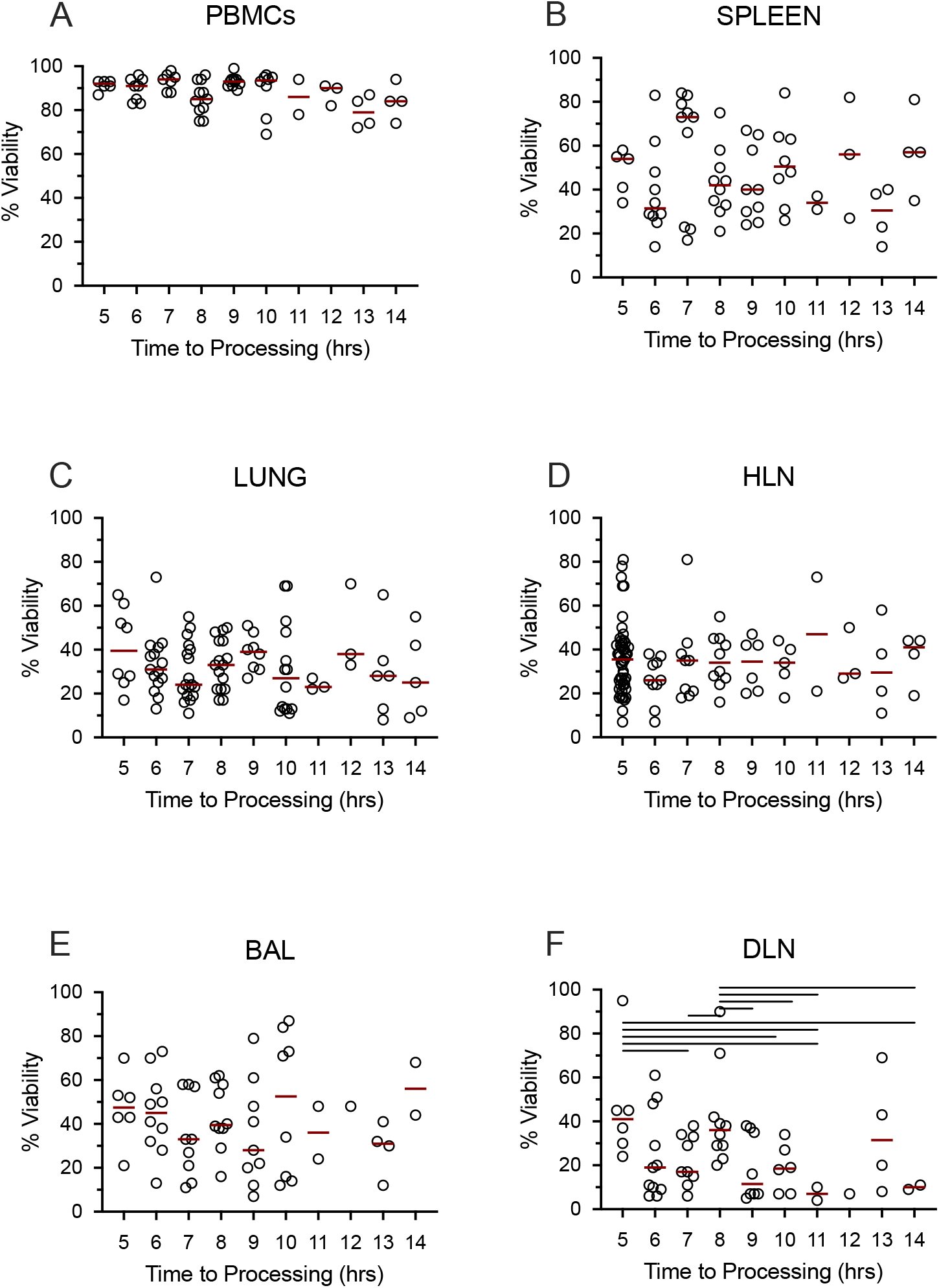
Tissue cells collected postmortem are viable up to 14 hours post death. Data shows percent viability of isolated single cells from the Blood (A), Spleen (B), Lungs (C), Hilar lung draining lymph nodes (D), Bronchoalveolar lavage (E) and distal, non-draining lymph node (iliac lymph nodes) (F). Lines above each graph represent a significant difference between the two groups at a significance level of p < 0.05. Data came from all subjects.

Having shown that we could isolate viable cells from the organs of deceased subjects, we next tested whether the cells were functional using the T-SPOT.TB assay (Figure 4). We found that PBMCs isolated postmortem were just as functional as PBMCs isolated from venous blood of a living subject (Figure 4A). Furthermore, the T-SPOT.TB assay worked on spleen and bronchoalveolar lavage (BAL) samples, showing that cells isolated from the tissues are also not just viable, but are functional as well. In undertaking the T-SPOT.TB assay, we were able to identify T-SPOT.TB negative and positive subjects (Figure 4B). It is important to note that data presented in Figure 4B are from deceased subjects from the S&E Unit. Figure 4B shows that 61% of the subjects tested were T-SPOT.TB positive – data very similar to other data in the same population (Biraro et al., 2014; Kizza et al., 2015).

**Figure 4.**
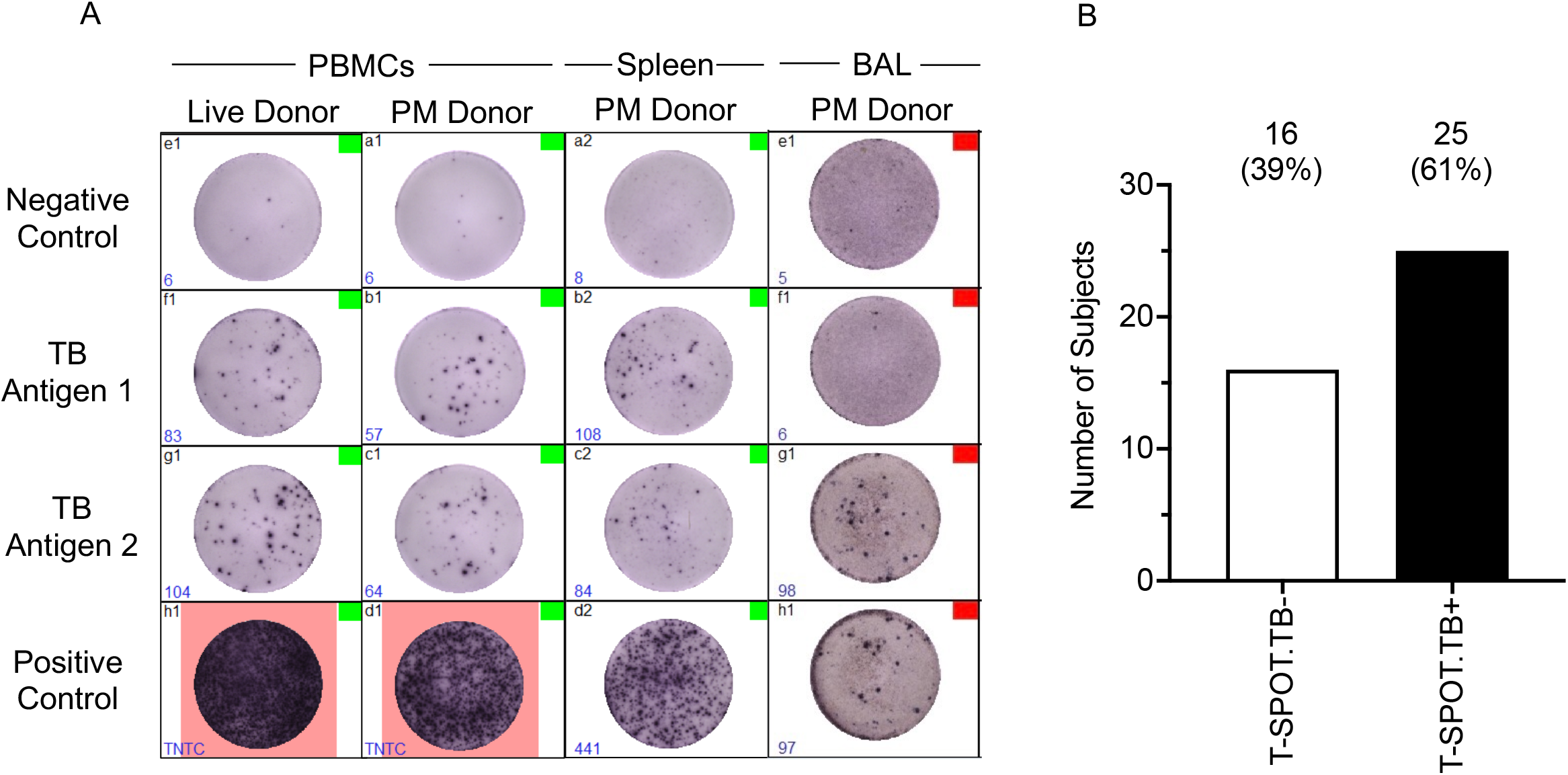
Tissue cells collected postmortem are functional in a T-SPOT.TB Assay. T-SPOT.TB data (Fig. 2A) shows that arterial PBMCs isolated and collected postmortem perform as well as venous blood from a live donor. In addition, the T-SPOT.TB assay also works in tissues such as the spleen and BAL. Culture conditions were slightly altered to enable the assay to work, as described in the methods. Fig. 2B shows latent TB infection (LTBI) data collected from our non-TB arterial blood samples, assayed using the T-SPOT.TB assay. The percentage T-SPOT.TB positive samples (25/16 – 61%), representative of a latent TB infection at the time of death, correlates closely with other published papers from this population [1, 2]. Data comes from HIV-/TB-people recruited from the SEU.

Next, we asked whether T cells isolated from different organs showed evidence of classical immunological naïve and memory cell subsets (Figure 5A and 5B; representative data). In all organs analysed, both CD4 (Figure 5A) and CD8 (Figure 5B) T cells had naïve (CCR7+/CD45RA+, T_N_), central memory (CCR7+/CD45RA-; T_CM_), effector memory (CCR7-/CD45RA-, T_EM_) and effector memory RA (T_EMRA_) subsets, albeit at different frequencies. Additionally, we also asked whether tissue resident memory (TRM) cells were present in the different organs, for both CD4 (Figure 5C) and CD8 (Figure 5D) T cells. Representative flow data are shown. Unsurprisingly, TRM cells (CD69+/CD103+) CD4 and CD8 T cells were absent in the blood (Figures 5C and D; PBMC), but present in all organs analysed (Figure 5C and D; LUNG, BAL and HLN).

**Figure 5.**
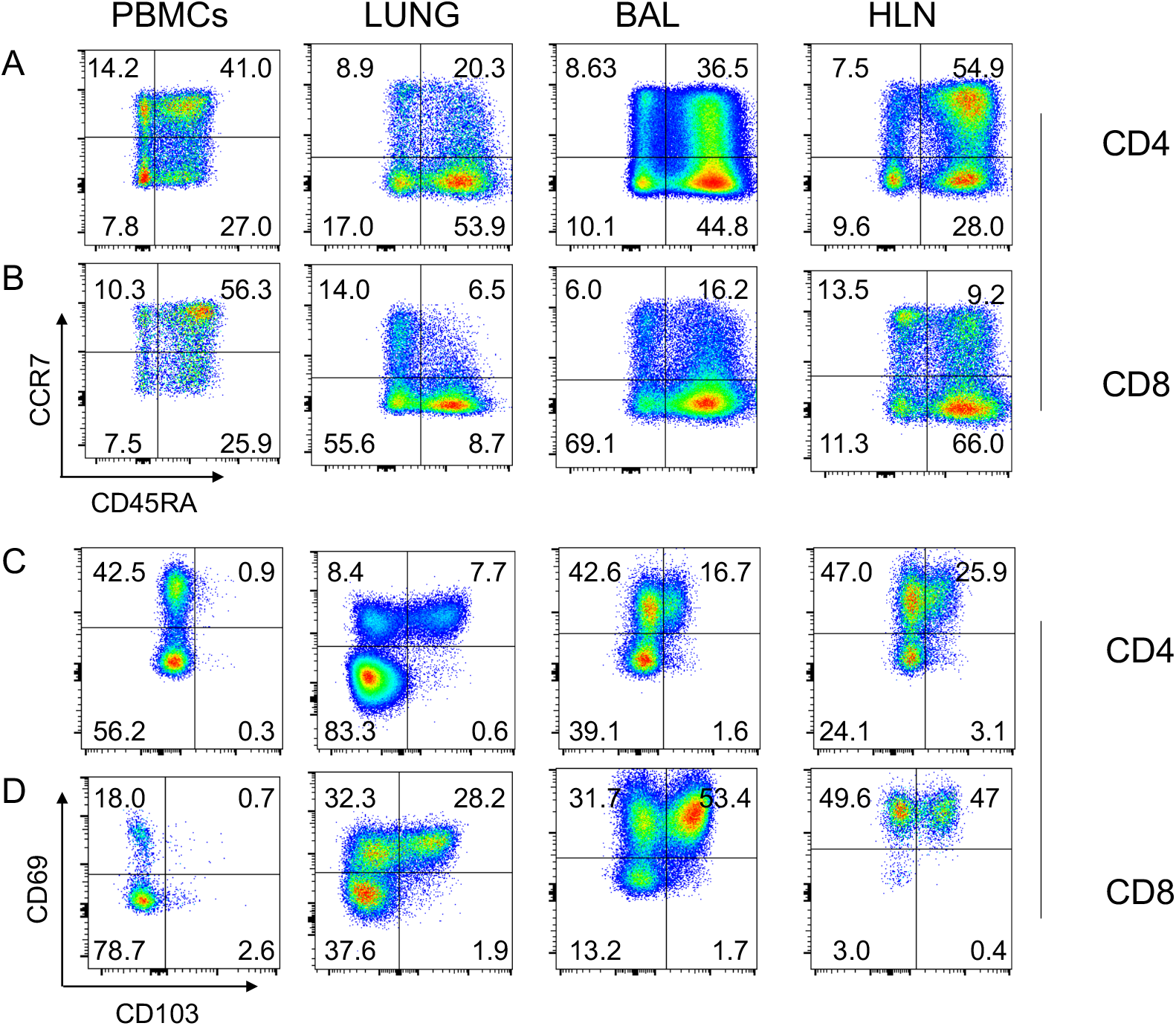
Naïve, memory and TRM subsets in different human tissues. Standard phenotypical markers (CCR7/CD45RA) were used to differentiate memory subsets within the indicated tissues for CD4 (Figure 3A) and CD8 (Figure 3B) T cells. Tissue-specific memory T cell markers (CD69/CD103) were also used to identify cell subsets within human tissue, again for both CD4 (Figure 3C) and CD8 (Figure 3D). Note that lung tissue-specific memory T cells (CD69+/CD103+) cells are absent from PBMCs, but present in lung tissue, BAL and HLN. Representative plots are shown, from an 74year old HIV-/TB-person recruited from the SEU. BAL, Bronchoalveolar lavage; HLN, draining lung hilar lymph node.

Finally, we asked which patient group (HIV+/PTB-, HIV-/PTB+, HIV+/PTB+ and disseminated TB (DISS TB)) stained positive for the different CCR7/CD45RA subsets in both CD4 (Figure 6A-D) and CD8 (Figure 7A-D) T cells. Note that all disseminated TB (DISS TB) subjects were HIV^+^ (data not shown). In general, there was no significant difference between any patient group and the CCR7/CD45RA subsets, except for the CCR7+/CD45RA+ cell subset within the BAL amongst CD4 T cells (Figure 6B, BAL), and the CCR7+/CD45RA+ and CCR7-/CD45RA-cell subsets amongst CD8 T cells (Figure 7B and D, respectively).

**Figure 6.**
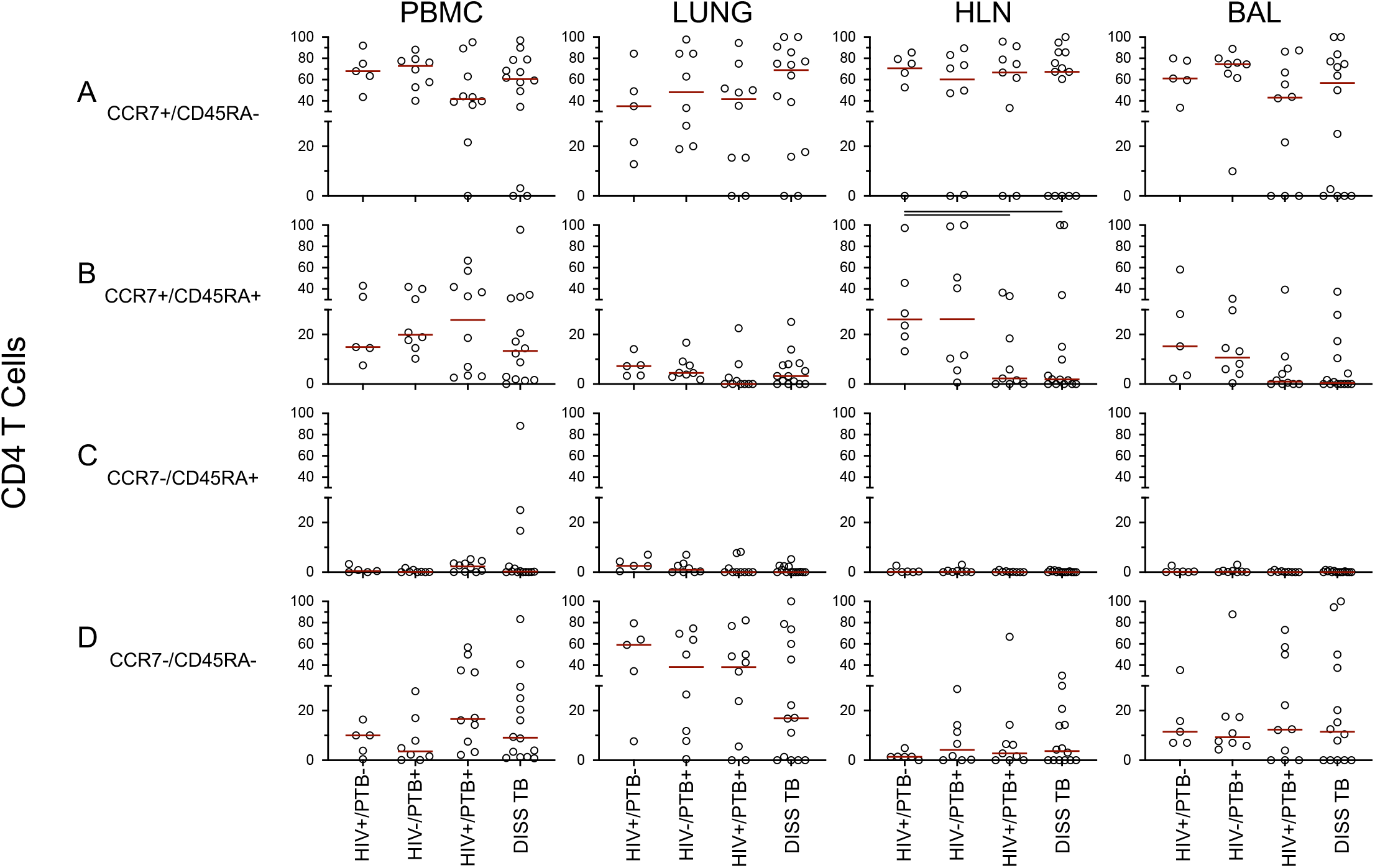
CD4 subset distribution within different clinical groups and organs based on a classical memory phenotype. Data shows results from the specified organs and clinical groups, based on classical memory (CCR7/CD45RA) phenotype. Data points represent results from a single individual. Bars above a plot represent a significant difference between the two clinical groups at a significance level of p < 0.05.

**Figure 7.**
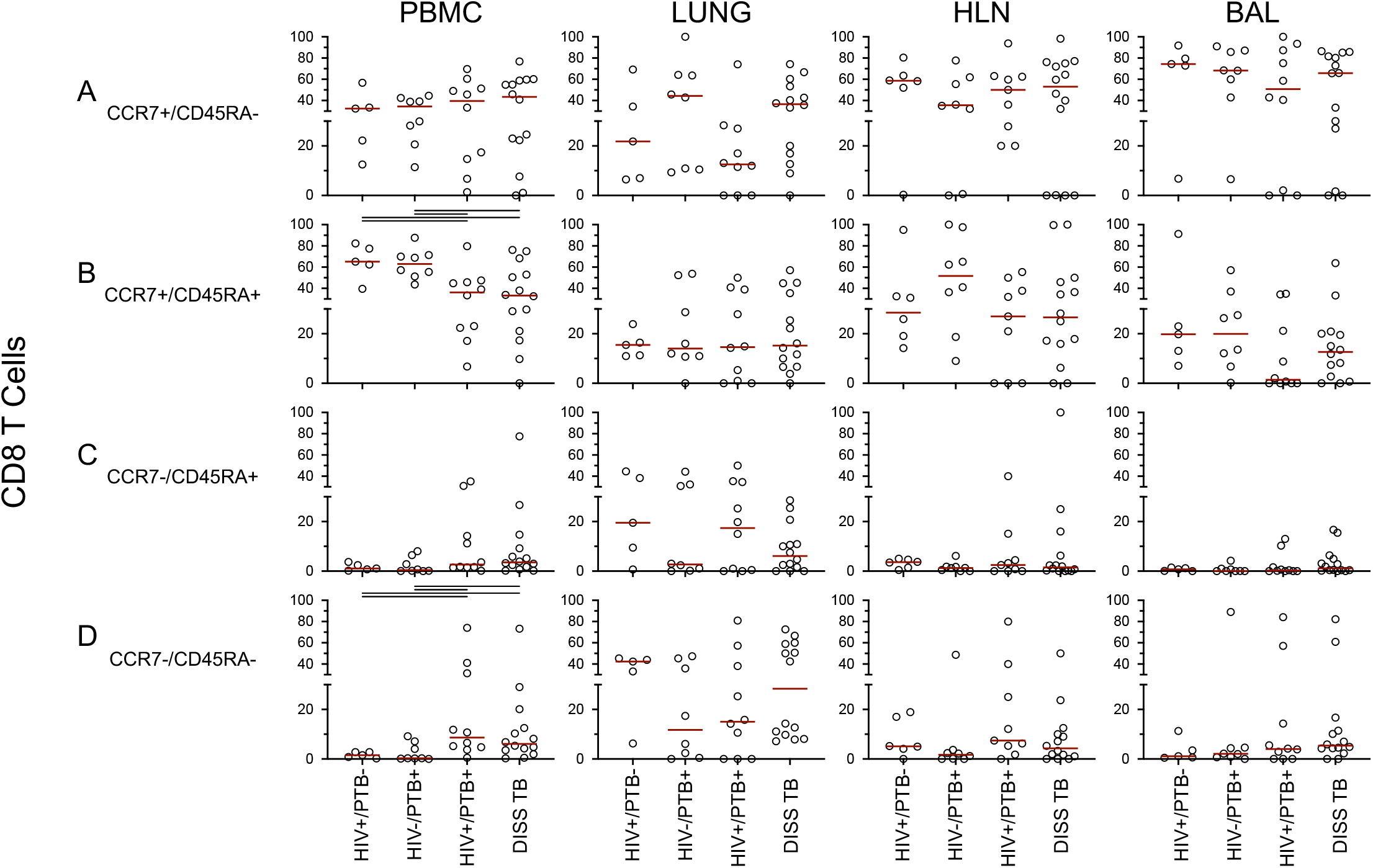
CD8 subset distribution within different clinical groups and organs based on a classical memory phenotype. Data shows results from the specified organs and clinical groups, based on classical memory (CCR7/CD45RA) phenotype. Data points represent results from a single individual. Data points represent results from a single individual. Bars above a plot represent a significant difference between the two clinical groups at a significance level of p < 0.05.

Similarly, we examined TRM cell subsets (CD69/CD103) amongst CD4 (Figure 8A-D) and CD8 (Figure 9A-D) T cells. Differences were observed between TB groups within the CD69^+^/ CD103^+^ and CD69^-^/CD103^+^ subsets for both CD4 (Figure 8B, C) and CD8 (Figure 9B, C) T cells in the lungs and BAL tissues. Double positive CD4 CD69^+^/CD103^+^ T cells were significantly lower in HIV+ subjects with or without TB by Mann Whitney tests, although this was lost after adjusting for multiple comparisons. However, the double positive CD8 CD69^+^/ CD103^+^ T cell subset (Figure 9B) in lungs were depleted in subjects that were TB and HIV co-infected. For BAL, the depletion of this subset was only seen in subjects that had disseminated TB (Figure 9B). These differences, however, were lost after correcting for multiple comparisons. There is some debate as to whether multiple testing is useful or not (Perneger, 1998). We present our data here with both multiple testing performed or not, for patient groups or cell subsets that show the greatest difference.

**Figure 8.**
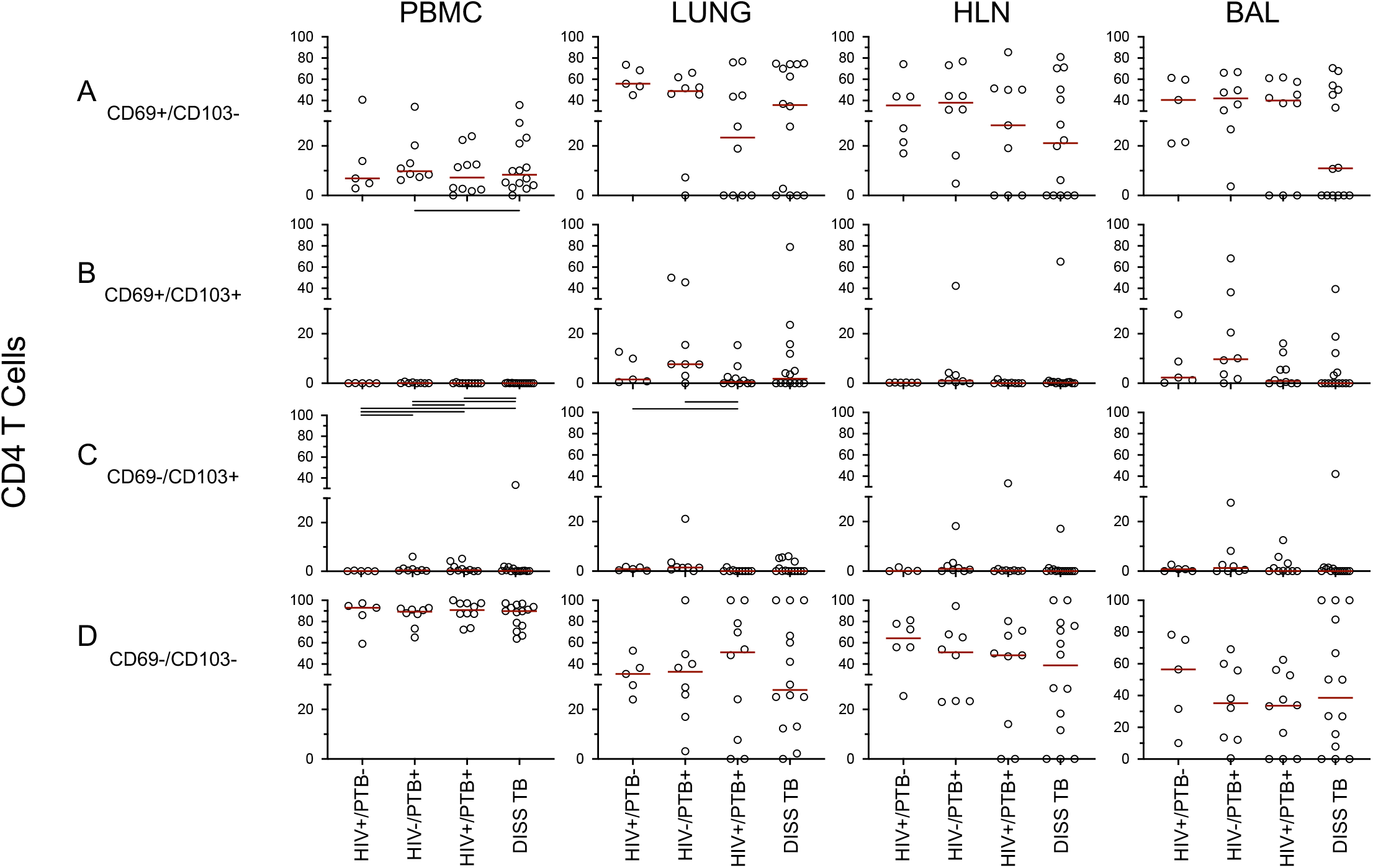
CD4 subset distribution within different clinical groups and organs based on a Tissue Resident Memory (TRM) phenotype. Data shows individual results from the specified organs and clinical groups, based on a TRM (CD69/CD103) phenotype. Data points represent results from a single individual. Data points represent results from a single individual. Bars above a plot represent a significant difference between the two clinical groups at a significance level of p < 0.05.

**Figure 9.**
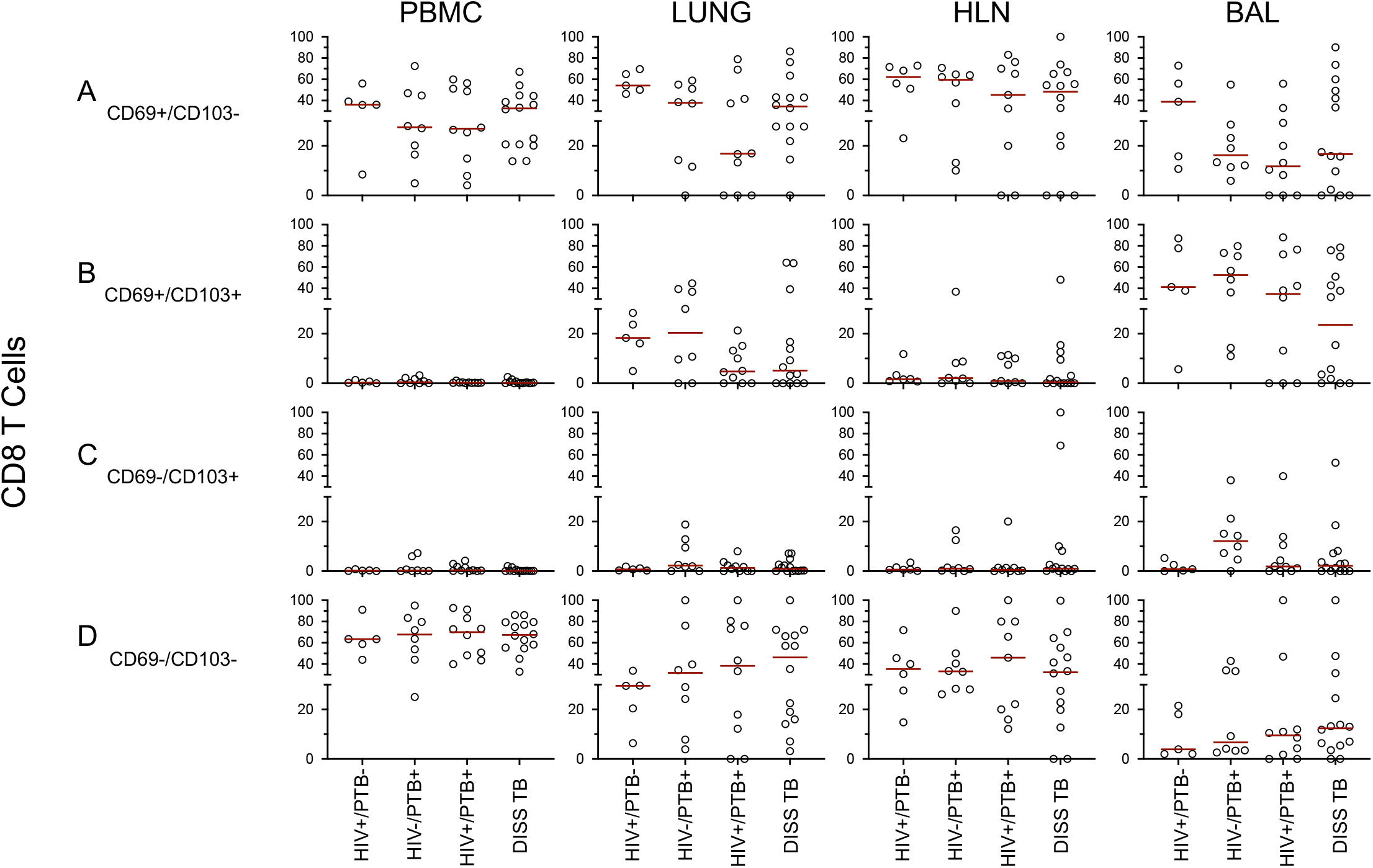
CD8 subset distribution within different clinical groups and organs based on a Tissue Resident Memory (TRM) phenotype. Data shows individual results from the specified organs and clinical groups, based on a TRM (CD69/CD103) phenotype. Data points represent results from a single individual.

Interestingly, the CD4 CD69^-^/CD103^+^ T cell (Figure 8C) depletion in the lungs of HIV^+^PTB^+^ subjects were picked by the Kruskal Wallis test. For the single positive CD8 CD69^-^/CD103^+^ T cells (Figure 9C) were depleted in HIV+ subjects with or without TB, going by Mann Whitney tests between PTB^+^HIV^-^ and either, PTB^-^HIV^+^, Disseminated TB or HIV^+^PTB^+^. These differences, however, were also not recognized by the Kruskal Wallis test of multiple comparisons. The CD4 and CD8 CD69+CD103-T cells (Figure 8A and 9A) were depleted only in the lungs of HIV^+^PTB^+^ subjects. It is important to note that the area of active TB infection in subjects with disseminated TB was the meninges. On the other hand, CD4 CD69 +CD103-T cells (Figure 8A) in BAL were depleted in subjects with disseminated TB, whereas their CD8 counterparts were depleted in any subject with TB, regardless of the clinical manifestation of TB, either with or without HIV.

## Discussion

Development of better TB vaccine strategies is based on deeper insights into the immunity underlying TB, and not just in blood, but most importantly at the site of infection. Most knowledge of TB immune responses at sites of infection is derived from animal models. However, translating results to humans has been limited by the inability to recapitulate the complex natural history of disease progression and reactivation as it exists in humans. Current human lung studies are sampled from patients with severe lung complications at the extreme end of the spectrum and lack normal lungs for comparison. In postmortem studies such as this, we are not only able to compare tissues of people who died of TB from those who died of other causes, but we can also study TB involved and other non-involved tissues within the same individuals. Unfortunately, postmortem studies have subsequently fallen out of favour for many reasons all over the world, but such studies are in understanding pathogen and immune system involvement at sites of infection. This TB postmortem study was set up to determine the feasibility of carrying out postmortem work for medical research in Uganda.

We report that coroner driven postmortems are feasible in Uganda. However, our findings show that this is highly dependent on whether the death is sudden or not. The 94% consent rate at the S&E ward is higher than what has previously been reported in Mulago National Referral Hospital (Cox et al., 2011), although this study looked across all wards at MNRH. Maixenchs and colleagues noted previously that circumstances such as sudden death were a major driving factor for willingness to have a postmortem undertaken (Maixenchs et al., 2016). Our data supports this finding; where most of the deaths are sudden (in our study, at the S&E Unit); the major reason given by the NOKs for postmortem consent was that knowing cause of death will help in coming to terms with their relative’s death. Indeed, the consent rate on the TB wards where death isn’t sudden was low. Additionally, the average length of hospital stay for TB cases was five days, during which a diagnosis had been established and therefore relatives had an idea of the probable cause of death and prognosis. Nevertheless, the consent rate on the TB wards was comparable to what has been observed previously in Mulago National Referral Hospital (Cox et al., 2011).

Phenotype, function, and location of immune cells within tissues is the basis of immunological research. To assess function, cells must be viable. Therefore, assessing viability of immune cells and whether sample integrity is maintained after death was a key objective for this study. Several studies have reported cell viability postmortem, with lymphocytes maintaining highest viability after death, and with viable cells identified in both blood and tissues after several days (Babapulle and Jayasundera, 1993; Dokgoz et al., 2001; Laiho and Penttila, 1981). Our average time from death to processing was nine hours (range 4hours to 14hours), and indeed, immune cells were still viable, functional and maintained sample integrity.

The main aim of this paper was to show that human postmortem studies are a valuable research tool to examine phenotype and function of cells within the tissue. We have shown that there are subtle differences in cell subsets between the blood (where most human research is focussed) and the tissues. TRMs play an essential role in containment of tissuespecific infections, and were present in tissues including the Lung, HLN and BAL, **but** absent within blood, as expected. Here we characterised the TRM CD4 and CD8 T cell profile based on CD69 and CD103 expression. Our data show gross changes rather than fine specificity or TB-specific populations within the different TB groups; further studies are ongoing to tease apart these differences.

HIV infection has previously been found to impair CD103 expression, which is important in the maintenance of cells in mucosal sites such as the lungs (Moylan et al., 2016). In agreement, the depletion of CD8 CD69^-^CD103^+^ T cells in BAL was associated with HIV. CD4 CD69^+^CD103^+^ depletion in BAL and lung was also associated with HIV. Interestingly, we found no depletion of the double positive CD8 CD69^+^CD103^+^ T cell subset in the lungs of either HIV+/TB- or HIV-/TB+ subjects but these cells were depleted in presence of both HIV and TB infections. This data suggests that the HIV associated T cell depletion is selective and only limited to double positive CD4 CD69^+^CD103^+^ subset in the lungs and BAL. Indeed, CD4 TRMs were shown to be primary targets of HIV infection (Cantero-Perez et al., 2019). More work needs to be done to understand the effect of HIV on the CD8 subsets.

We report that the CD4 CD69^+^CD103^-^ in BAL was the only cell subset associated with TB disease only. However, depletion of both CD8 and CD4 CD69^+^CD103^-^ subsets in the lungs of subjects with PTB+HIV+ co-infection, but not in subjects with disseminated TB (all disseminated TB subjects had HIV and little or no involvement of the lungs), suggesting that the CD69^+^ activation which has been seen in HIV and TB alone (Moylan et al., 2016; Yang et al., 2020) doesn’t span to the lungs when there is a co-infection of both TB and HIV. Taken together, our results highlight complexity of the relationship and interactions between *M.tb*, HIV and host immune response.

Examining TB-specific cell subsets within the tissues is a major priority of our future research. We expect significant differences to occur between antigen-specific T cell subsets within the organs, and most notably within the lung, granuloma and draining lymph nodes. Further research is required to understand these subsets during health and disease.

## Conclusion

We have shown in this study that postmortem studies are a feasible approach to study tissue-specific diseases. Lymphocytes can be isolated from tissues up to 14 hours after death, with no loss of viability. In addition, cells are functional and show tissue specific surface markers. We did not measure antigen-specific T cell responses other than undertaking a T-SPOT.TB assay. Further research in this area is sorely needed, not just for understanding TB disease and its progression, but any disease with a tissue-specific tropism. Although this is missing in this study, we nevertheless show that postmortem procedures remain a valuable and essential tool to both establish cause of death, and to advance our medical and scientific understanding of disease.

## Acknowledgements

This work was supported by NIH Contract 75N93019C00070 and was conducted at the MRC/UVRI and LSHTM Uganda Research Unit which is jointly funded by the UK Medical Research Council (MRC) part of UK Research and Innovation (UKRI) and the UK Foreign, Commonwealth and Development Office (FCDO) under the MRC/FCDO Concordat agreement and is also part of the EDCTP2 programme supported by the European Union. We are grateful to the administrations of Mulago National Referral Hospital and Kiruddu National Referral Hospital for their enthusiasm and cooperation in this research. We extend our utmost gratitude to the bereaved families who gave permission to enrol deceased relatives into the study. We also acknowledge the outstanding efforts of the clinical team who made this study possible.

